# A sensory ecological perspective on mate sampling strategies: simulation models and an empirical test

**DOI:** 10.1101/533489

**Authors:** Diptarup Nandi, Megha Suswaram, Rohini Balakrishnan

## Abstract

Long-range communication signals play a central role in mate search and mate choice across a wide range of taxa. Among the different aspects of mate choice, the strategy an individual employ to search for potential mates (mate sampling) has been less explored despite its significance. Although analytical models of mate sampling have demonstrated significant differences in individual fitness returns for different sampling strategies, these models have rarely incorporated relevant information on the ecology of signalers and sensory physiology of receivers, both of which can profoundly influence which sampling strategy is optimal. In this study, we used simulation models to compare the costs and benefits of different female mate sampling strategies in an acoustically communicating field cricket (*Plebeiogryllus guttiventris*) by incorporating information on relative spacing of callers in natural choruses, their signal intensity and the effect of signal intensity on female phonotaxis behaviour. Mating with the louder caller that the female first approaches emerged as the optimal strategy, thus reflecting the importance of physiological mechanisms of sound signal localization (passive attraction) over active sampling. When tested empirically in the field, female behaviour was consistent with passive attraction.

## Introduction

Searching for mates is a critical component of animal mating behaviour. Mate sampling strategies are employed by individuals of the choosier sex (typically females) in their search for potential mates (Andersson 1994). In systems where sexual selection acts via the mechanism of mate choice, active mate sampling strategies can directly influence the strength of sexual selection (Andersson 1994; Reinhold and Schielzeth 2014). The kind of sampling strategy employed and the number of potential mates sampled influence the extent to which mate choice affects the evolution of sexual traits (Gibson and Langen 1996; Roff and Fairbairn 2015). Despite the importance of sampling strategies for understanding sexual selection, empirical studies have largely focused on measuring sexual trait variation, choosiness for these traits and mating preferences (Reinhold and Schielzeth 2014).

A number of theoretical studies have, however, investigated mate sampling strategies utilizing analytical models (Janetos 1980; Parker 1982; Wiegmann et al. 1996; Johnstone 1997; Kokko et al. 2014). In the threshold or sequential sampling strategies, females evaluate mates sequentially based on a certain threshold value of the trait and decide to mate only if the mate quality is higher than the threshold criterion that either remains constant or declines with increasing costs of sampling (Janetos 1980; Real 1990; Dombrovsky and Perrin 1994; Wiegmann et al. 1996). In the other category of models, mating decisions are based on a comparative assessment of the previously sampled potential mates; either between the last pair of successively sampled individuals (‘Sequential-comparison’), or a fixed number of sampled individuals (‘best-of-n’) (Janetos 1980; Wittenberger 1983). Unlike ‘threshold’ strategies, the ‘comparison’ strategies allow for a potential mate to be revisited during sampling and hence require the female to remember the location and/or the identity of a previously sampled individual (Janetos 1980). Models with and without search costs yielded maximum fitness returns for the ‘One-step decision’ and the ‘Best-of-n’ strategy strategies respectively (Janetos 1980; Real 1990). More recent models have employed information theoretic approaches to reduce the uncertainty of information gathered on mate quality by allowing for multiple revisits to potential mates (‘Bayes tactic’, “threshold” strategy and ‘Random walk’: Luttbeg 1996; Wiegmann and Angeloni 2007; Wiegmann et al. 2010; Castellano and Cermelli 2011).

A critical feature of all active mate sampling strategies is the ability of females to reject mates or defer their mating decision until further sampling (Parker 1982). In mating systems in which females localize mates using their sexual advertisement signals, the potential mate with a more intense advertisement signal can, however, attract more females due to a larger broadcast area (Parker 1982, 1983). This phenomenon, referred to as ‘passive attraction’, may superficially appear like active female choice (Parker 1983; Kotiaho and Puurtinen 2007). However, a female’s attraction in this case is guided entirely by its physiological response to the relative strengths of the perceived advertisement signal and does not involve active mate sampling (Parker 1983). A distinction between active mate sampling strategy and ‘passive attraction’ is critical especially because female ‘choosiness’, a component of mate choice, has been defined as the extent of mate sampling that it engages in before mating (Kokko et al. 2006; Kotiaho and Puurtinen 2007).

Empirical studies on mate sampling strategies demonstrate varied strategies being used by animals belonging to different taxa, suggesting a role of the ecology of the system in determining the optimal strategy. For instance, while some species of birds have been reported to use either ‘best-of-n’ or the ‘Bayes tactic’ (Trail and Adams 1989; Dale et al. 1990, 1992; Petrie et al. 1991; Bensch and Hasselquist 1992; Hovi and Rätti 1994; Rintamäki et al. 1995; Luttbeg 1996; Uy et al. 2001), others have been found to employ ‘sequential comparison’ and ‘one step decision’ strategies (Choudhury and Black 1993). Studies on mate sampling strategies in fishes and invertebrates have generally provided evidence on the use of sequential threshold strategies (Bakker and Milinski 1991; Milinski and Bakker 1992; Forsgren 1997; Reid and Stamps 1997).

During mate search animals often utilize communication signals wherein, typically, individuals of one sex use signals produced by the other sex to localize and evaluate mates. Consequently, in long-range communication for mate attraction, the ecology of signalers and the sensory physiology of receivers can profoundly affect mate sampling strategies. The extent of time spent signaling and movement during signaling can constrain the time window of female mate search, since some of the ‘comparison-based’ strategies require the female to revisit an already sampled male. Relative spacing of signalers determines the amount of movement involved in female mate search. Therefore, the optimal mate sampling strategy will vary depending upon the energetic and predation costs of female mate search, and the relative spacing of males. Moreover, signal intensity determines the extent of broadcast of a signal and its effect on receiver response to the signal can be critical (Parker 1983; Kotiaho and Puurtinen 2007). The number of signalers that a female can simultaneously perceive is determined by the degree of overlap between the broadcast areas, which in turn depends on the relative signal intensities and spacing of signaling males (Forrest and Raspet 1994; Murphy and Gerhardt 2002). Neither theoretical models nor empirical research of mate sampling strategies, however, have incorporated relevant sensory ecological information of the communication system.

Lack of simultaneous availability of information on the ecology of signalers in wild populations and the sensory physiology of responders has severely constrained our understanding of the ecological effects on mate sampling strategies. Elaborate experimentation on the auditory sensory physiology of acoustic signal reception and detailed characterization of structurally simple acoustic signals in orthopterans and anurans provide potential model systems to explore the aforementioned questions (Gerhardt and Huber 2002). Empirical studies on female mate sampling strategies in some species of crickets have however been conducted under laboratory conditions mainly using playback experiments, rendering them difficult to interpret in a natural context since rejection of loudspeakers may not imply rejection of calling males (Wiegmann 2000; Wagner Jr. et al. 2001; Beckers and Wagner Jr. 2011). More conclusive are field studies in anurans, where females were observed to mate with the first encountered male closest to the female position (Murphy and Gerhardt 2002; Meuche et al. 2013). Since the closest males are also likely to be the loudest at the female position, a lack of measurement of signal intensity in these studies limits the interpretation of the role of acoustic signals in female mate sampling. Song intensity is particularly pertinent because it is well established to play a critical role in female phonotactic response, and can override female preferences for other call features (Stout and McGhee 1988; Castellano et al. 2000; Nandi and Balakrishnan 2013).

Previous studies on the ecology of callers and the effect of sensory physiological mechanisms on female phonotaxis behaviour, proffers the field cricket species *Plebeiogryllus guttiventris* as a suitable model system for exploring female mate sampling (Mhatre and Balakrishnan 2006; Nandi and Balakrishnan 2013). *P. guttiventris* males can form dense choruses where individual callers maintain their calling site within a night (Mhatre and Balakrishnan 2006), but move frequently across multiple nights of calling, with inconsistent calling activity across the calling nights (Nandi and Balakrishnan 2016). Moreover, callers are also consistent in the different acoustic features of their calling songs within a night of calling activity (Nandi and Balakrishnan 2013). These facts suggest that the time window available to females for sampling mates is more likely to be limited to the calling period of males within a night. Results based on playback experiments in this species further demonstrate that the calling song sound pressure level (SPL), a measure of signal intensity, profoundly affects female phonotaxis behaviour and overrides female preference for faster calls (Mhatre and Balakrishnan 2007; Nandi and Balakrishnan 2013). Females approach the sound source that is louder at the female position and not at source (Mhatre and Balakrishnan 2008). Moreover, females do not approach the closest sound source, demonstrating the dominant role of signal intensity over source proximity in female evaluation of mates (Mhatre and Balakrishnan 2008). Models based purely on the auditory physiology of females were able to accurately predict phonotactic trajectories of female approach, further illustrating the dominant role of calling song SPL in female phonotaxis behaviour (Mhatre and Balakrishnan 2007, 2008).

Despite the considerable impact played by signal intensity across all the different sensory modalities of communication, studies on mate sampling strategies have seldom investigated their effects (Kotiaho and Puurtinen 2007). According to the ‘passive attraction’ model, signal intensity directly affects the mating success of the signaler by attracting more females (Parker 1983). However, the likelihood of localizing high intensity signalers will depend on whether a female evaluates signal intensity at the female position or at source and also on the extent of overlap between the broadcast ranges of the signalers. Searchers may also increase their fitness by mating with more intense signalers if signal intensity indicates male quality. For instance, in *P. guttiventris* male calling song SPL in *P. guttiventris* has high repeatability and is the only calling song feature that correlates positively with male body size (Nandi and Balakrishnan 2013). Therefore searchers may adopt active sampling strategies that maximize the likelihood of mating with high intensity signalers, to overcome the constraints imposed by their sensory physiology and the ecology of the communication system.

In this study, we investigated female mate sampling strategies, with a combination of simulations based on natural distributions of male calling SPL and relative spacing, and experiments on calling males and searching females in their natural habitat, using *P. guttiventris* as a model system. In the simulations, we compared costs and benefits of using passive attraction with those of active sampling strategies, under the sensory ecological constraints of the communication system. In the experiments we specifically asked whether females reject males based on their calling song SPL at source in order to infer the possible mate sampling strategy that females use.

## Methods

### Field site and animals

All field work was conducted in privately-owned agricultural fields near the village Ullodu, in the Chikballapur district of the southern Indian state of Karnataka (13°38′48.81″ N, 77°42′45.23″ E), with consent and permission of the owners. In the period between February and May from 2010 to 2013, all the observations and experiments were conducted between 1900 and 2200 hours, the peak calling period of male *P. guttiventris*. These were either collected from the field population as nymphs close to their last instar (n =14) or were from a laboratory culture (n =19) that has been maintained since 2008 and outbred with individuals from the wild population at regular intervals of approximately a year (Nandi and Balakrishnan 2013).

### Male spacing and calling song SPL

Data on spatial location of males and their calling song SPL measurements were collected for a total of 11 choruses in the field. Choruses were identified by acoustic sampling of the natural habitat, tracking individual callers and confirming calling behaviour of males visually. Habitat patches were specifically selected where male callers aggregate spatially for many nights. All calling males within a chorus were identified and their calling sites were marked with caller ID-annotated flags. Calling song SPL of all the callers was measured at 20 cm in front of the caller (referred to as calling song SPL at source) at an angle of 90° with respect to the raised wings, using a Brüel & Kjær ½” microphone, Type 4189 (20 Hz to 20 kHz) and a Sound Level Meter, Type 2250 (Brüel & Kjær, Naerum, Denmark) set at fast root mean squared (RMS) time weighting. Distances of the marked calling sites from a few centrally located reference points and the angles subtended were measured using a meter tape and survey precision compass mounted on a tripod (Survey Compass 17475780, error ±0.5°, conceptualized by Francis Barker and Sons Ltd., sold and serviced by Lawrence and Mayo, India). These distances and angles were used to generate the Cartesian co-ordinates of individual calling sites for all the choruses using custom-written scripts in Matlab version 7.11.0 (R2010B) (Math-Works, Natick, MA, U.S.A.).

### Mate sampling simulations: General

Simulations based on caller spacing in natural choruses, corresponding call SPL and female phonotactic rules, were used to compare the performance of different mate sampling strategies. All the simulations were performed in Matlab version 7.11.0 (R2010B) (Math-Works, Natick, MA, U.S.A.). Calling male spacing data from 11 natural choruses were used to generate chorus maps containing the Cartesian co-ordinates of the callers and the corresponding call SPLs were used generate their broadcast area (area around a calling male where the sound signal is audible to a female). The broadcast area was estimated as a circle with the caller at the center and the radius being determined by the distance at which the calling song SPL attenuates to the mean female hearing threshold (40 dB SPL re 2 × 10^−5^ N/m^2^) in this species (Mhatre and Balakrishnan 2006).

At the initiation of a simulation, virtual females were assigned positions randomly within a circular area, with a radius of 20 m that was derived from the summation of the maximum distance of callers from a central reference point across all the eleven choruses and the maximum broadcast radius of the callers. The area of this circle, encompassing all the callers within a chorus with their acoustic broadcast ranges, was used as the sampling space for females (Fig. S1). Once assigned a position, it was assessed whether the female could hear any of the callers by comparing the linear distances (*d*_*fm*_) between the female position and the callers with that of their respective broadcast radii (*br*_*m*_). If the linear distances were greater than the broadcast radii of all the callers (*d*_*fmi*_ > *br*_*mi*_ for all *i*), then the female was considered to be outside the broadcast area of any caller. If *d*_*fmi*_ < *br*_*mi*_ then the female could hear the *i*^*th*^ male.

A female followed a random walk outside the signal broadcast area with a speed that varied stochastically for every step within a run, with a mean movement of 0.1 m/minute and a standard deviation of 40 percent of the mean. After every five seconds, the female position was assessed to check if it lay within the broadcast area of any caller as described above. If the female was within the broadcast area of a male, she approached that male (center of the circle) with a speed of 0.07 m/minute (based on the mean time taken by a female to reach a male in the field phonotaxis experiment).

A female could hear multiple males at certain spatial co-ordinates where the broadcast areas of two callers overlapped. In such scenarios, a female approached the male that was louder at its position if the difference in SPLs between the callers was more than 3 dB at the point of evaluation, or else the female chose randomly between the callers that it could hear. The evaluation of louder callers was repeated four times while the female was on its path towards the louder male to ensure that females had more than one opportunity to evaluate the louder caller.

### Mate sampling simulations: Strategies

A female was given a maximum of 180 minutes (3 hours) to sample males which was based on previous observation of peak calling activity in wild populations of *P. guttiventris* and low across-night calling site fidelity (Nandi 2016). In those 180 minutes a female evaluated male calls every five seconds, leading to 2160 evaluation steps. After localizing a male, a female’s mating decision was dependent on the sampling strategy that it employed. In the simulations females were tested on five different sampling strategies:

#### ‘Random Sampling’ (RS)

A female mated with the first male encountered (spatial proximity of less than 5 cm with the male) while moving randomly. In this strategy, females did not evaluate males based on their calling song and hence did not show phonotaxis and consequently the outcome is essentially based on random movement of the females.

#### ‘Fixed Threshold’ (FT)

Females sampled males based on their calling song SPL and mated with a male only if its calling song SPL was greater than the threshold value, which was set at the mean (77.2 dB) of the SPL distribution measured previously for this *P. guttiventris* population (Nandi and Balakrishnan 2013). If the calling song SPL was less than the threshold, the female continued its mate search either until it found a caller with calling song SPL above the threshold or exhausted the maximum sampling time.

#### ‘*One-step decision*’ (OS)

This strategy was similar to the ‘fixed threshold’ strategy, except the threshold declined as a function of increasing costs (time of searching). The calling song SPL threshold (*I*_*th*_) was assumed to be a function of time spent searching (*t*) given by the equation *I*_*th*_ = *a* + *be*^−*ct*^ where *a*, *b* and *c* are constants. The starting threshold (at *t* = 0) was chosen to be the maximum SPL for the range of calling song SPLs pooled across all the 11 choruses, instead of the population mean to discount for the subsequent decay in mate quality (Real 1990). The slope of the curve was selected such that the threshold declined steeply to the mean SPL value after sampling for 22% of the time window available for sampling (*t* = 0.22 × 180) to ensure minimal rejection of mates based on the higher thresholds. The curve was made to asymptote at the minimum value of the SPL range.

#### ‘*Best-of-n*’ (BN)

Females sampled a maximum of 5 males or for 180 minutes and mated with the male with the highest calling song SPL at source. In this strategy a female was allowed to mate only if it sampled more than one male (n > 1). Females mating with a male after having sampled just that male (n=1), is similar to ‘passive attraction’. The minimum requirement of sampling more than one male in case of ‘best-of-n’ was therefore necessary to distinguish it from ‘passive attraction’.

#### ‘*Passive attraction*’ (PA)

Females mated with the first male that they localized based on the calling song SPL at female position and never rejected any male.

The number of females for a given run of the simulation was kept the same as the number of calling males in the chorus (sex ratio = 1:1), for all the five sampling strategies. The entire process was iterated a hundred times for each of the five sampling strategies in 11 choruses. In every iteration, the initial co-ordinates of the females were varied randomly.

Benefits to females employing each of the sampling strategies were assessed in terms of the probability of mating and the probability of mating with a louder male. The probability of mating was calculated by dividing the number of females that mated by the total number of females used in the simulations using a particular strategy within a chorus. The probability of mating with a louder male was calculated by dividing the number of females that mated only with males which were louder than mean calling song SPL by the total number of females. To compare the costs to females using different sampling strategies, sampling time, defined as the time expended by a female in finding a mate, was calculated for every female that found a mate in each of the iterations. The number of males sampled before choosing a mate was also calculated for each of the females that found a mate following the three active sampling strategies (FT, OS, BN).

### Mate sampling simulations: exploring the input parameter space

The initial conditions and values of different parameters in the simulation could affect the relative performance of the females using different mate sampling strategies. Increasing the rate of movement of females and time window available for sampling and decreasing the area of sampling can enhance a female’s likelihood of encountering potential mates. Since different sampling strategies depend variably on the encounter rates, the relative performance of the sampling strategies may vary in consequence. Therefore, time available for sampling, rate of female movement and the area of sampling were varied systematically to investigate the effect of these input parameters. The simulations described above were repeated, increasing the sampling time available for females from the initial 3 hours (180 minutes) to 6 hours (180×2 minutes) and 9 hours (180×3 minutes), separately. Simulations were also run for each of these three sets, with a higher rate of female movement outside the broadcast areas with a mean of 1 m/minute and a standard deviation of 40 percent of the mean. Therefore, there were two sets with different rates of female movement, ‘slow females’ (0.1 m/minute) and ‘fast females’ (1 m/minute). The six sets of simulations (3 time windows for sampling × 2 rates of female movement) were repeated with a reduced area of sampling. In the ‘Small’ area sets, circles were constructed for each of the choruses separately, with radii estimated as the sum of the maximum distance of a caller from the centroid of the chorus and its corresponding broadcast radius for a given chorus (range of area reduction: 10-95%, Fig. S1). Therefore, the simulations were conducted separately for a total of 12 parameter combinations.

### Mate sampling simulations: Statistical analyses

The main objective of our analyses was to compare and contrast the benefits and costs between females using ‘passive attraction’ and the active sampling strategies. P-values estimated under the frame-work of ‘null hypothesis tests’ can lead to biased interpretations when the sample sizes are very high particularly in the context of simulations (White et al. 2014). Instead, estimations of effect sizes and confidence intervals have been argued as a more appropriate alternative to ‘null hypothesis testing’ (Nakagawa and Cuthill 2007) and were consequently used to interpret our simulation results. All statistical analyses were conducted using R version 3.3 (R Development Core Team 2014). The effect sizes for the probability of mating and the probability of mating with a louder male were estimated as the median of the bootstrapped pairwise differences between the probabilities for females using ‘PA’ strategy and other active sampling strategies over 1000 iterations, for every chorus separately. The effect sizes for the sampling time were estimated as the mean difference between the bootstrapped means of sampling times across choruses for females using ‘PA’ strategy and other strategies. 95 percent confidence intervals were estimated using the bootstrapped distributions with 1000 iterations. Confidence intervals not overlapping zero imply a less than 5 percent likelihood of observing the difference by chance alone.

### Mate sampling experiment

This experiment was designed to test whether female crickets actively reject some callers in favor of others and if so, to distinguish between the mate sampling strategies in use. The experimental design had to ensure that the female crickets be presented with the best chance of rejecting a male i.e. scenarios where the females were most likely to approach a male with a suboptimal trait value while simultaneously perceiving the presence of another male. Pairs of male *P. guttiventris* calling males were located in the field such that the distance between them was less than a meter to ensure female’s perception of both males simultaneously (Mhatre and Balakrishnan 2006). After measuring calling song SPL at source, a pair was selected for the experimental trial only if one of the callers called at an SPL less than the mean SPL (77.2 dB) of the population except in 4 trials (where both were higher). If multiple pairs of callers were located within a night, the pair with the least distance between the two callers or maximum relative SPL difference at source was selected for experimentation. The relative SPL of male calling song at the female release position was calculated using the known attenuation profile of *P. guttiventris* calling song in the habitat (Mhatre and Balakrishnan 2006). Two CCTV cameras were deployed to monitor the calling activity of males throughout the duration of the experiment, which were connected to a laptop via a DVR system placed at a distance of more than 7 m from the callers. The experiment was initiated 5-10 minutes after setting up the CCTV cameras to ensure minimal disruption in calling activity.

Virgin females, aged 10 to 13 days after the final molt (see Supplementary Material for details), were released approximately 40 cm from the caller with the lower calling song SPL at source (1^st^ male) such that the release point was collinear with the two callers and the 1^st^ male was louder at the female position than the 2^nd^ male (louder at source). Once released, the female was followed using either an IR-sensitive video-camera or by eye under red light from a distance to ensure minimal disruption of its movement. Female approach to a calling male was confirmed by two observers, one directly observing the females and the other observing CCTV transmission of male activity on the laptop. Female mating was confirmed either by female mounting behaviour followed by male-female coupling and subsequent transfer of the male spermatophore, when the mating behaviour took place outside the cracks in the ground from which the males were found calling, or by relocating the female with a spermatophore attached when matings took place inside the cracks. The experimental observation was continued until the unmated male stopped calling or the mated female walked away from the calling males. In trials where there was no mating after phonotaxis, observations were continued until the males stopped calling or the female walked away from the chorus.

### Remating experiment

Laboratory experiments were conducted on *Plebeiogryllus guttiventris* females to investigate the probability of remating and the time taken since the first mating to do phonotaxis and remate. In the first set, single-speaker playback and mating trials were conducted on the same females for at least five consecutive nights after the first mating. In an independent second set, females were tested in the same experimental paradigm with a latency of a week from the first mating for two weeks. Laboratory bred females between the age of 11-17 days since the final moult were used for these experiments. All experimental trials were conducted inside an anechoic chamber. Phonotactic and mating behaviors were recorded using an IR sensitive video camera (Sony, DCR TRV 17-E, Sony Corporation, Japan) under external infrared lights.

A female was released 50 cm from a speaker that played out a typical *P. guttiventris* call with mean song parameter values appropriate for that temperature (25° C). The SPL at the female release position was ensured to be at 61 dB (typical SPL value at 50 cm from the source in the field) using a Brüel & Kjær Sound Level Meter, Type 2250 with a ½” microphone, Type 4189. Females were presented with a conspecific male after showing a phonotaxis response, defined as reaching within 5 cm from the speaker or within 10 minutes from the initiation of the trial when phonotaxis response was absent. The mating trials were conducted in a sponge cube of 35.5 cm x 35.5 cm x 30 cm with a cylindrical hole of diameter 15 cm and depth 15 cm in the center. A mating trial was scored as negative only if the male courted and the female did not mount the male after 10 minutes from the start of the mating trial. A female was exposed to a new male for the subsequent trials. Identical experimental methodology was followed for the two different sets.

### Ethical Note

All the SPL measurements, spatial data collection and animal handling for the behavioral experiments were as per the national guidelines for the ethical treatment of animals.

## Results

### Mate sampling simulations

The choruses sampled in the field varied in the number of callers (4-13) and nearest neighbour distances between callers (Fig. S2). The effect sizes for the probability of mating was significantly greater than zero for all the comparisons across the different simulation sets and therefore implies that the females using ‘passive attraction’ were the most likely to mate (Fig. 1). Despite low magnitude of effect sizes, the percentage change was considerable even for comparisons with the fixed-threshold strategy (ranging from 15.6 % to 37.9 %) which yielded the least difference (Fig. 1 and Fig. S3). The relative performance of different sampling strategies did not change across the different treatments of time window for sampling and sampling area within the ‘Fast’ and ‘Slow’ female sets (Fig. S3).

**Figure 1.**
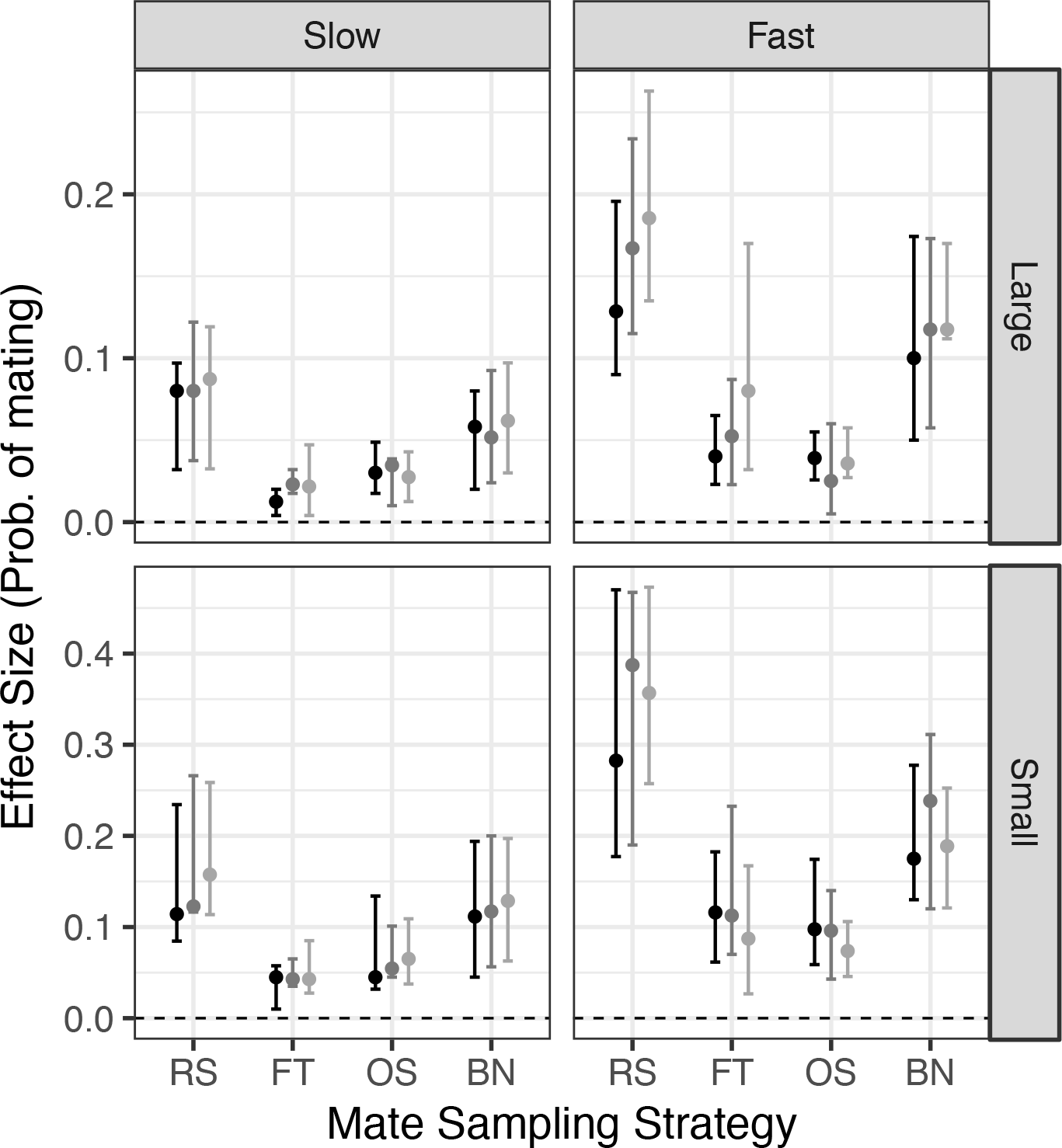
Effect sizes of the probability of mating. The circles and the error-bars represent the medians and confidence intervals of the bootstrapped distributions of the pairwise differences (choruswise) in the probability of mating between ‘passive attraction’ and the other sampling strategies. Black = 3 hours, Grey = 6 hours, Light grey = 9 hours time window available for sampling. RS = Random sampling; FT = Fixed-threshold; OS = One-step decision, BN= Best-of-n. ‘Large’ = Large Area, ‘Small’= Small Area.

The probability of mating with a louder male for females using the ‘passive attraction’ strategy did not significantly differ from those using the active sampling strategies (Fig. 2). Only the ‘Fast’ females with 9 hours sampling time window in ‘Large’ area using ‘Fixed-threshold’ strategy yielded significantly higher values than ‘passive attraction’ (Fig. 2). Comparisons with the ‘best-of-n’ strategy yielded significantly positive effect sizes only in 2 out of 12 scenarios, despite higher magnitudes in the rest of the sets (Fig. 2 and Fig. S4).

**Figure 2.**
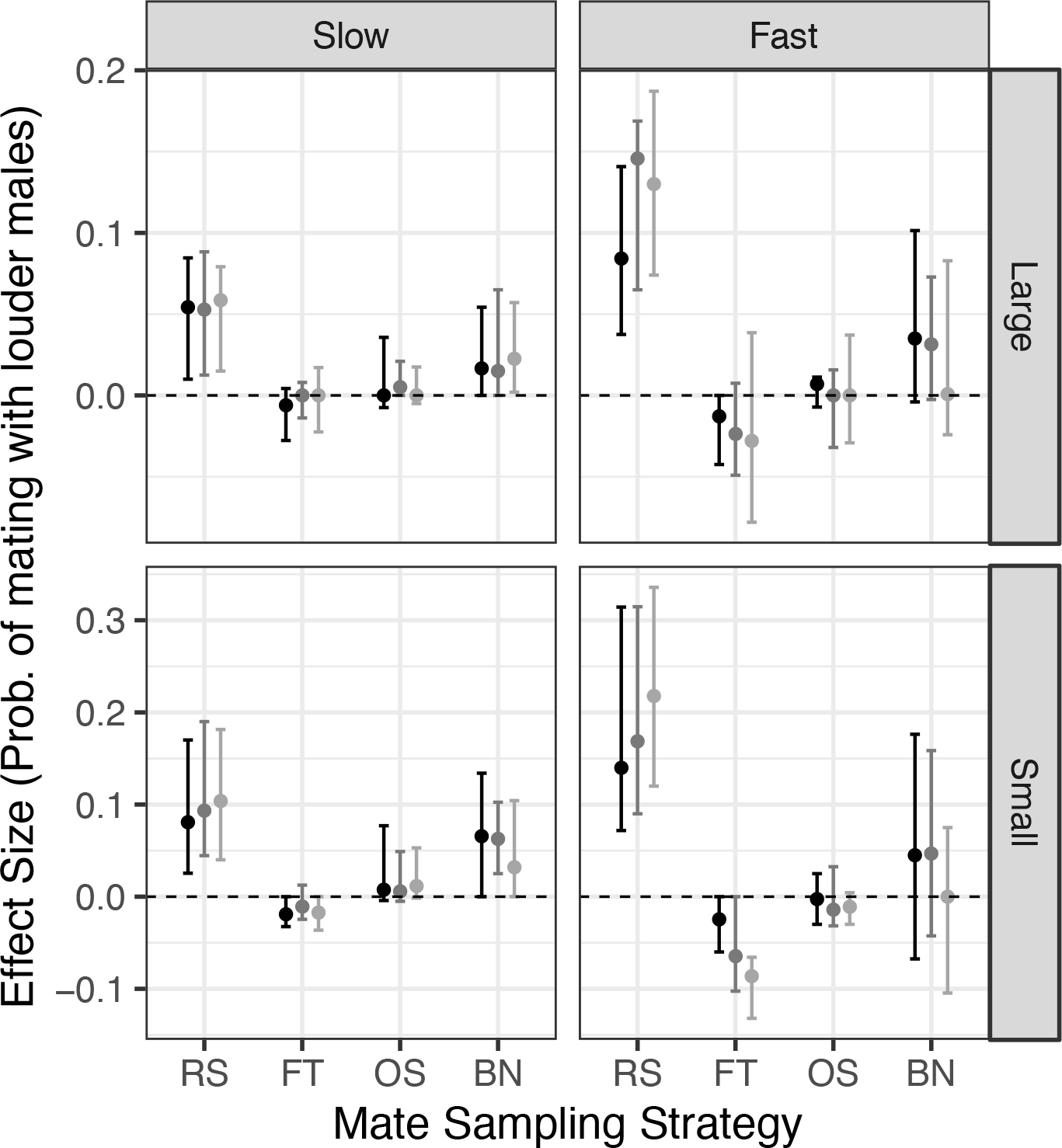
Effect sizes of the probability of mating with a louder male. The circles and the error-bars represent the medians and confidence intervals of the bootstrapped distributions of the median pairwise differences (choruswise) in the probability of mating with louder males between ‘passive attraction’ and the other sampling strategies. Black = 3 hours, Grey = 6 hours, Light grey = 9 hours time window available for sampling. RS = Random sampling; FT = Fixed-threshold; OS = One-step decision, BN= Best-of-n. ‘Large’ = Large Area, ‘Small’= Small Area.

Females using the ‘best-of-n’ strategy utilized the entire time window available for sampling, consequently spending the maximum time searching for mates across all the sets (Fig. S5). In absence of acoustic cues, females that searched for mates randomly (“RS”) also spent more time searching for mates compared to other strategies (Fig. S5). Effect sizes significantly lower than zero in comparisons between the ‘passive attraction’ and the threshold strategies (‘FS’ and ‘OS’) in most of the simulation sets implies that females spent the least time searching for mates while using ‘passive attraction’ (Fig. 3). Only in 2 out of 12 scenarios, the time spent sampling for females using the ‘fixed-threshold’ strategy did not significantly differ from that of ‘passive attraction’ (Fig. 3). Females using ‘passive attraction’ could reduce their sampling time by a minimum of 10 % when compared to the threshold strategies. Most of the ‘Slow’ and ‘Fast’ females (> 75%) sampled only one male before mating while using ‘fixed-threshold’ and ‘one-step decision’ strategies (Fig. 4 and Fig. S6). Females using the ‘best-of-n’ strategy could sample more males with increasing the rate of movement, time window for sampling and in the smaller sampling area (Fig. S6).

**Figure 3.**
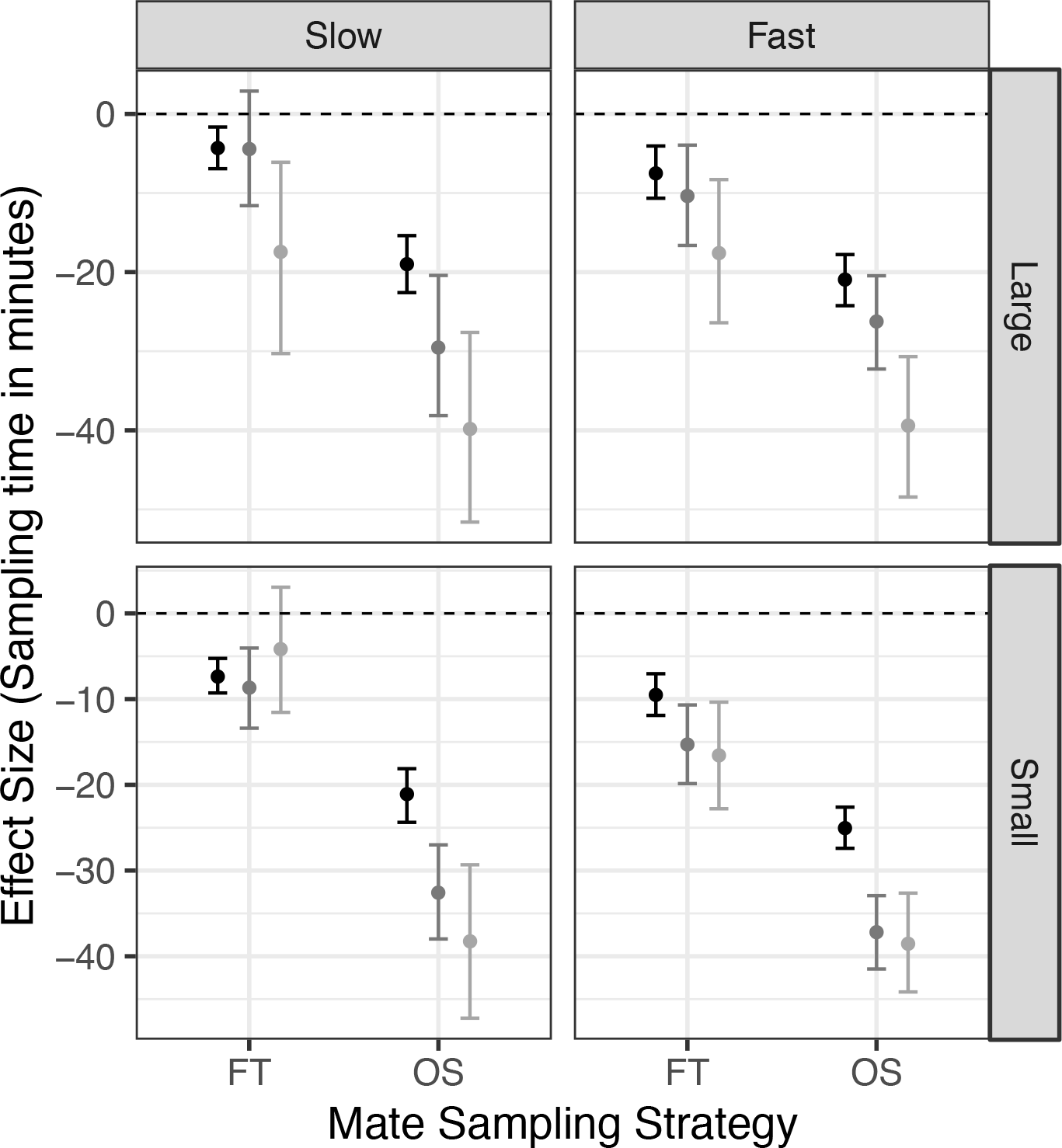
Effect sizes of the time spent sampling for females that mated. The circles and the error-bars represent the means and confidence intervals of the distributions of the mean differences in the time spent sampling between ‘passive attraction’ and the threshold strategies, as estimated by bootstrap sampling. FT = Fixed-threshold; OS = One-step decision. ‘Large’ = Large Area, ‘Small’= Small Area.

**Figure 4:**
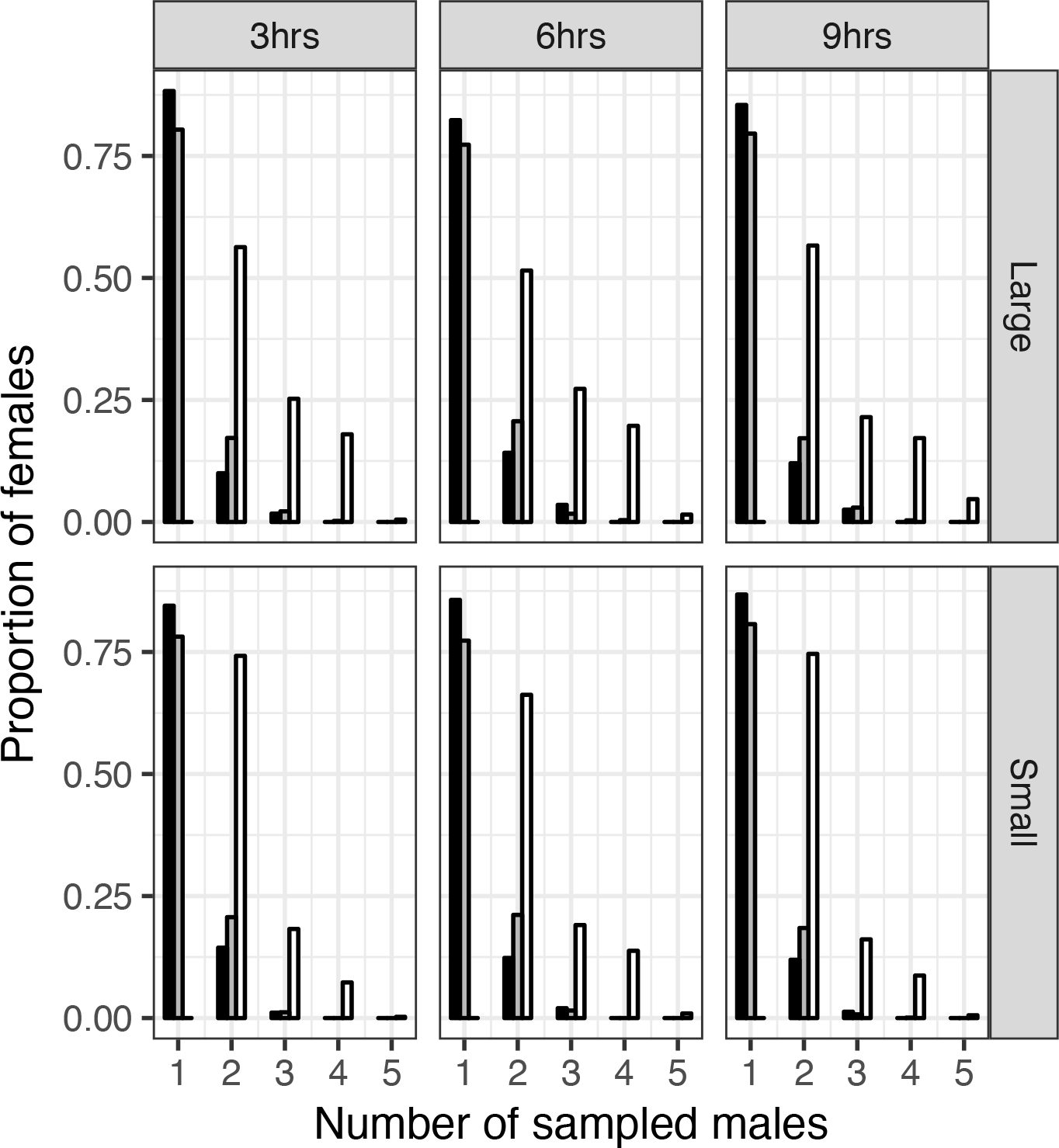
Proportion of sampled males. Relative frequency distribution of the number of males sampled by females that were successful in mating, using each of the three active sampling strategies for ‘Slow’ females. Black = 3 hours, Grey = 6 hours, White = 9 hours of time window available for sampling.

The probability of mating while using ‘passive attraction’ was higher than both the ‘fixed-threshold’ and ‘one-step decision’ strategies despite lowering the thresholds (Fig. S7a). The magnitude of the difference (effect size) however diminished on reducing the threshold. The difference in the probabilities of mating with a louder male between ‘passive attraction’ and ‘fixed-threshold’ strategies remained insignificant even after lowering the threshold (Fig. S7b). However, on reducing the starting threshold, ‘Slow’ females using ‘one-step decision’ strategy were more likely to mate with a louder male than females using ‘Passive Attraction’ (Fig. S7b).

### Mate sampling experiment

Ten out of 33 trials were dropped because in five of them the females did not show phonotaxis, in two trials the second male had stopped calling by the time of first mating and in three trials matings could not be confirmed. In the remaining 23 trials, a significant majority (20) of the females mated with a male after showing phonotaxis (Chi square test: *χ*^2^ = 12.56, *P* = 0.0004). None of the females approached the second male after localizing the first male, even in cases where they did not mate with the first male (3/23 trials). Thus, females predominantly mated with the first male they localized and never approached the second caller. Both the laboratory bred (*n* = 11) and wild caught females (*n* = 9) followed the same strategy of mating with the first male approached irrespective of their nymphal growth conditions.

Out of the 20 trials in which matings were observed, all females mated with the first male approached despite their calling at a lower SPL (at source) than the second male (Fig. 5a) (*Exact binomial test*: *P* < 0.0001). Only in two out of 20 trials was the male that the female mated with the one that called at an SPL higher than the population mean of the SPL distribution (Fig. 5a) (*Exact binomial test*: *P* = 0.0004). In ten out of the 18 trials where the first male called at an SPL below the population mean, the calling song SPL was even lower than one standard deviation below the mean (Fig. 5a). Both the males were audible to females in all the 20 trials as their calling song SPL at the female position was higher than the phonotactic hearing threshold of 40 dB (Fig. 5b). Moreover, the first male was louder than the second male at the female release position in all the trials (Fig. 5b).

**Figure 5:**
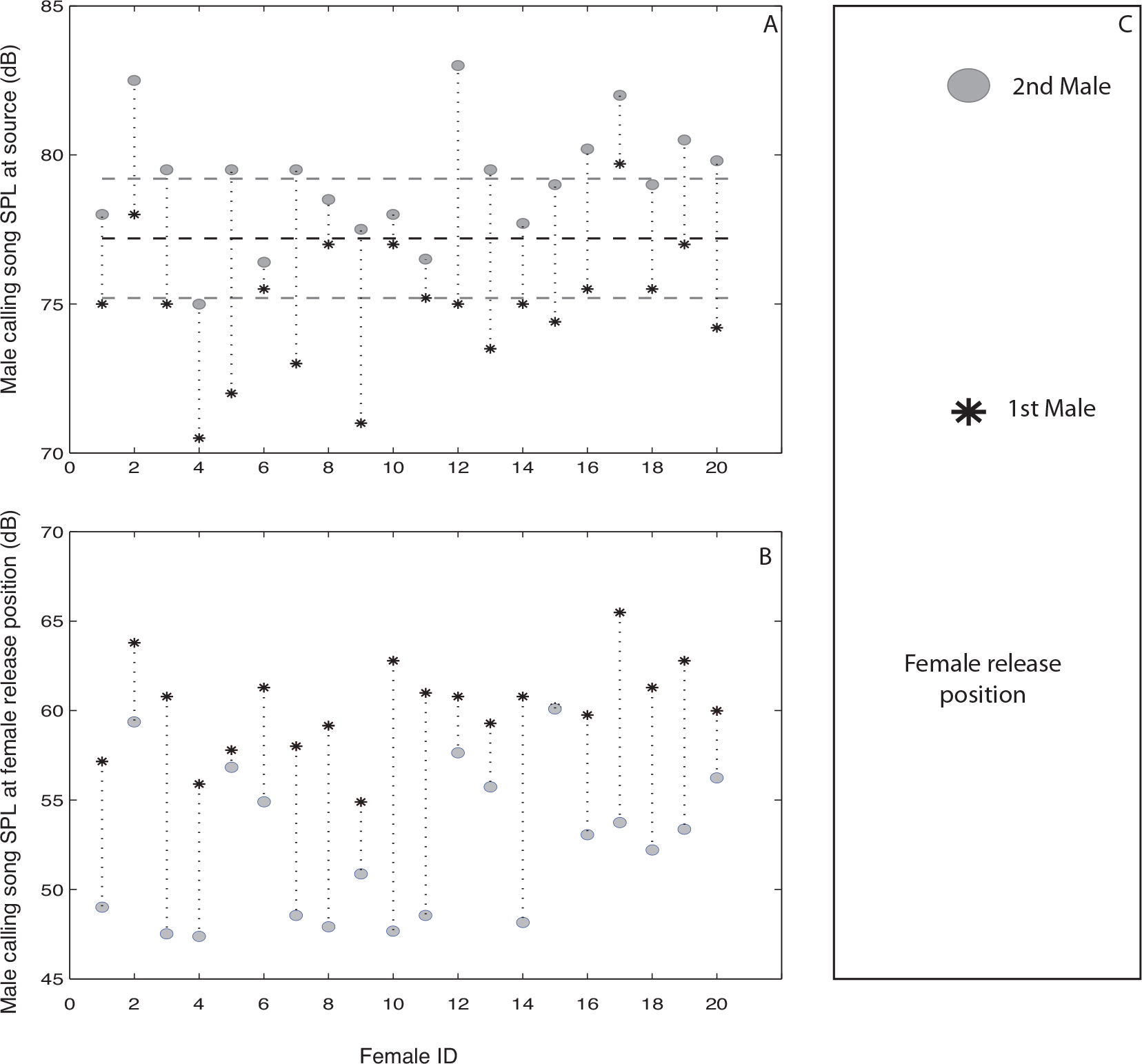
Female mating in relation to male calling song SPL. Calling song SPL of the pair of males per trial (A) at source and (B) at the female release position, in each of the 20 trials where matings were observed. Within a trial, the 1^st^ male is denoted by a star and the 2^nd^ male by a filled circle. The mated males are denoted by an asterix. The pair of males tested in a trial is joined by a vertical dotted line. The horizontal black dashed line marks the mean call SPL of the population with the standard deviation on either side represented by gray dashed lines. (C) A diagrammatic representation illustrating the positions of first and second males relative to the female release position

### Remating experiment

In the first set, out of the 27 females that mated as virgins, 11 (40.7 %) remated at least once when tested on six consecutive days. In the second set, out of 21 females that mated as virgins, 11 (52.4 %) remated at least once when tested on two consecutive weeks. The modal value of latency between the first and the second mating (refractory period) across days was one day (Table S1). In the second set, females remated after a week’s refractory period (Table S1). Only 1 (9.1 %) and 2 (18.2 %) out of independent sets of 11 mated females showed phonotaxis before remating. Therefore, the probability of showing phonotaxis prior to remating was significantly lower than that of a virgin and does not increase with increasing time since the last mating (1/11 mated versus 14/27 virgin in the first set: *χ*^2^ = 4.33, *P* = 0.038; 2/11 mated versus 14/21 virgin in the second set: *χ*^2^ = 4.99, *P* = 0.026).

## Discussion

### ‘Passive attraction’ as an optimal search strategy: simulations

Among all the different sets of simulations, ‘Slow’ females with a sampling window of 3 hours in a ‘Large’ area of sampling is the ecologically most relevant scenario in the case of our study system, the field cricket *P. guttiventris*. Females responding to an acoustic signal in the field moved at a speed comparable with the ‘Slow’ females. Hence, females are unlikely to move faster outside the signal broadcast area unless they fly (which is rarely observed). Calling activity within a night is usually limited to a peak period in most acoustically communicating species and callers may change their calling site across multiple nights as has been demonstrated in certain species of crickets (Ritz and Köhler 2007; Nandi and Balakrishnan 2016). In such scenarios the sampling period of females is likely to be limited by the calling activity period within a night.

In the ecologically relevant simulation set (‘Slow’ females with 3 hours sampling window in ‘Large’ area), females using ‘passive attraction’ were most likely to mate. The probability of mating depends on the rate of encounters between the searching female and the signaling male. Performance of active sampling strategies depends more heavily on encounter rates as they rely on sampling more males. However, even after relaxing the values of parameters such as rate of female movement, sampling window and sampling area, to increase encounter rates, females using ‘passive attraction’ performed significantly better than other active sampling strategies in terms of mating probability. Moreover, even after a reduction of the threshold criterion for ‘fixed-threshold’ and ‘one-step decision’ strategy, females following those strategies failed to match the performance of ‘passive attraction’

While probability of mating indicates the efficiency of finding mates, probability of mating with louder males indicates fitness benefits, as call SPL is assumed to be correlated with fitness (Nandi and Balakrishnan 2013). By definition, active sampling strategies are expected to yield higher probability of mating with louder males, due to the threshold criterion for selection or comparison of SPLs at source (Janetos 1980; Real 1990; Gibson and Langen 1996). In the simulations, however, active sampling strategies could not outcompete ‘passive attraction’ with respect to mating with a louder male. Increasing the encounter rates by relaxing the values of the input parameters, such as ‘Fast’ females or longer sampling durations (9 hours) or lower initial threshold, did lead to an enhanced performance of active sampling strategies but still failed to perform better than ‘passive attraction’ in most cases. In burrowing field crickets, callers are known to maintain their calling site across multiple nights and therefore allow females to sample for longer durations (Forrest and Green 1991; Rodríguez-Muñoz et al. 2010, 2011). Dense choruses can also provide a smaller area of sampling. However, our simulations demonstrate that, even after reducing the ecological constraints, active sampling strategies based on acoustic cues rarely outperform ‘passive attraction’ in finding a mate with higher fitness.

In the simulations, among the active sampling strategies, ‘best-of-n’ was severely affected by the ecological constraints. Females using the ‘best-of-n’ strategy could not sample more than 4 males, the optimal *n*, due to the constrained time window available for sampling and restricted female movement relative to the spatial separation between callers. The suboptimal sampling opportunity led to diminished performance. This result mirrors the consequences of incorporating costs into the analytical models of active mate sampling, leading to suboptimal performance of the ‘best-of-n’ sampling strategy (Real 1990; Dombrovsky and Perrin 1994; Wiegmann et al. 1996).

Females that succeeded in mating using ‘passive attraction’ took the least time to find mates since any encounter with a mate always guaranteed a mating, unlike in active sampling strategies. Time optimization can be critical to a sampling strategy, considering the possible costs such as energetic expenditure, time investment and increased exposure to predation (Real 1990). Search costs have been empirically shown to play an important role in mate sampling behaviour in both vertebrates and invertebrates (Milinski and Bakker 1992; Wickman and Jansson 1997; Byers et al. 2005; Kasumovic et al. 2007; Berger-Tal and Lubin 2011). The costs of mate searching could thus lead to the use of a strategy that not only maximizes the probability of finding mates with better quality, but also reduces the time spent searching.

In the simulations, females using either an active sampling strategy or ‘passive attraction’ always approached a caller guided by their auditory physiology. Furthermore, females using the ‘fixed-threshold’ and ‘one-step decision’ strategies were as likely to mate with a louder male as while using ‘passive attraction’, thus suggesting a mechanistic similarity. This similarity is corroborated by the fact that even in the threshold strategies, females predominantly mated with the first approached male, thereby reducing the requirement of their behavioural rules. Thus our simulations demonstrate that the physiological rule of approaching males louder at the female position can be sufficiently beneficial to females without employing active mate sampling strategies. Therefore, females using ‘passive attraction’ based purely on well-established physiological rules of sound localization (Mhatre & Balakrishnan, 2008) performed optimally across all the different sets of the simulations.

### Female mate sampling strategies: experimental results

The fact that females approached only one of the males in all experimental trials and never rejected a male in favour of the other male, indicates that they were not using a ‘best-of-n’ strategy, which requires a certain number of males to be sampled before choosing a mate (Janetos 1980; Real 1990; Gibson and Langen 1996; Wiegmann et al. 1996). If females were using male calling song SPL at source as thresholds to evaluate potential mates, then mated males should have called at an SPL higher than the population mean (Janetos 1980; Real 1990). However, 90% of the males that the females mated with called at SPLs lower than the mean population SPL and 50% of the males called at SPLs even lower than one standard deviation below the mean SPL. Thus, virgin females of *P. guttiventris* are unlikely to be using either of the threshold strategies. The results of our experiments instead suggest that female mating decisions are dictated by evaluation of male calling song SPL at female position and not at source, that is, females are using the ‘passive attraction’ strategy. A critical distinction between active mate sampling strategies and ‘passive attraction’ is the rejection of some potential mates over others (Parker 1982, 1983). In this study, a small proportion (13.6%) of females did not mate with the first male they approached, but, importantly, these females never approached the second male. Theoretical models of female phonotaxis as well as empirical studies of phonotaxis, particularly in this species, have demonstrated the importance of relative SPL difference at the female position in determining female approach to sound sources (crickets and anurans: Forrest and Green 1991; Forrest and Raspet 1994; *P. guttiventris*: Mhatre and Balakrishnan 2007, 2008). In this study we show that virgin females not only localize males that are louder at the female position but also mate with them, and do not actively sample calling males before deciding to mate.

Mating status could potentially affect the mate sampling strategy employed by females by making them choosier after the first mating. A mated female would invest in mate search only when the benefits of a second mating exceed the costs incurred by searching for males. In the laboratory experiments, *P. guttiventris* females remated, at times, when males were offered in close proximity, but rarely responded phonotactically. This implies that mated females, despite their motivation to remate, are rarely using male acoustic signals to search for potential mates. Moreover, latencies after mating, which were more than sufficient to induce remating, did not increase the likelihood of phonotaxis. The lack of phonotaxis prior to mating therefore raises pertinent questions regarding the importance of male calling song for mated females. In a few pilot trials (n =5) of the mate sampling experiment in the field with mated females, females did not approach either of the callers, despite being tested after the latency period from their first mating. Our results, therefore, suggest that mated females are unlikely to use a mate sampling strategy based on acoustic sampling of potential mates and points to differences in mating strategies between virgin and mated females that merit further investigation.

## Conclusion

A simulation framework incorporating information on natural distributions of calling song SPL and male spacing, and the physiology of female phonotaxis behavior, enabled an ecologically relevant comparison of female sampling strategies. ‘Passive attraction’ to the advertisement signal emerged as an optimal solution to mate search when ecological factors such as male spacing, female movement and total time available for sampling constrained sampling strategies. Moreover, in experiments using calling males in the wild, we demonstrate that virgin *P. guttiventris* females use passive attraction rather than an active sampling strategy to search for mates. ‘Passive attraction’ could possibly be a general female mating strategy in acoustically communicating animals considering the sensory ecological constraints of the communication system (Murphy and Gerhardt 2002; Meuche et al. 2013). Future studies on mate sampling, both theoretical and experimental, should therefore consider incorporating information on the sensory physiology of receivers and the ecology of signalers.

## Appendix

## Materials

### Animal maintenance

The collected nymphs were maintained individually in plastic boxes (14 ×10×5 cm) with moistened cotton wads and fed *ad libitum* with juvenile dog food (Pedigree, Mars India Pvt Ltd.) and Calcium Sandoz (Novartis India Ltd.). The date of their final moult was noted down and the adult was maintained in the same container till the end of the experimental trial. For the set from the culture, male and female nymphs were maintained separately in 50-litre plastic barrels. Female nymphs were transferred to similar plastic boxes as described above on the day of their moulting into adults. All the females were marked on their pronotum with unique colour codes using a nontoxic paint marker (Edding 780, Edding, St Albans, U.K.) for individual identification.

### Varying threshold parameters in the simulations

A lowering of the threshold SPL for females following the threshold strategies could increase the probability of mating for active sampling strategies relative to passive attraction. To investigate the effect of varying thresholds on the performance of females using threshold strategies, two sets of simulations were conducted separately. In the first set, the threshold criterion for females using the fixed-threshold strategy was reduced from mean SPL to 1 SD below the mean SPL for simulations with both ‘Fast’ and ‘Slow’ females with 3 hours time window for sampling in the ‘Large’ sampling area. In the second set, the initial threshold for females using one-step decision strategy was reduced to the mean SPL for simulations with both ‘Fast’ and ‘Slow’ females with 3 hours time window for sampling and in the ‘Small’ area of sampling.

**Figure A1.**
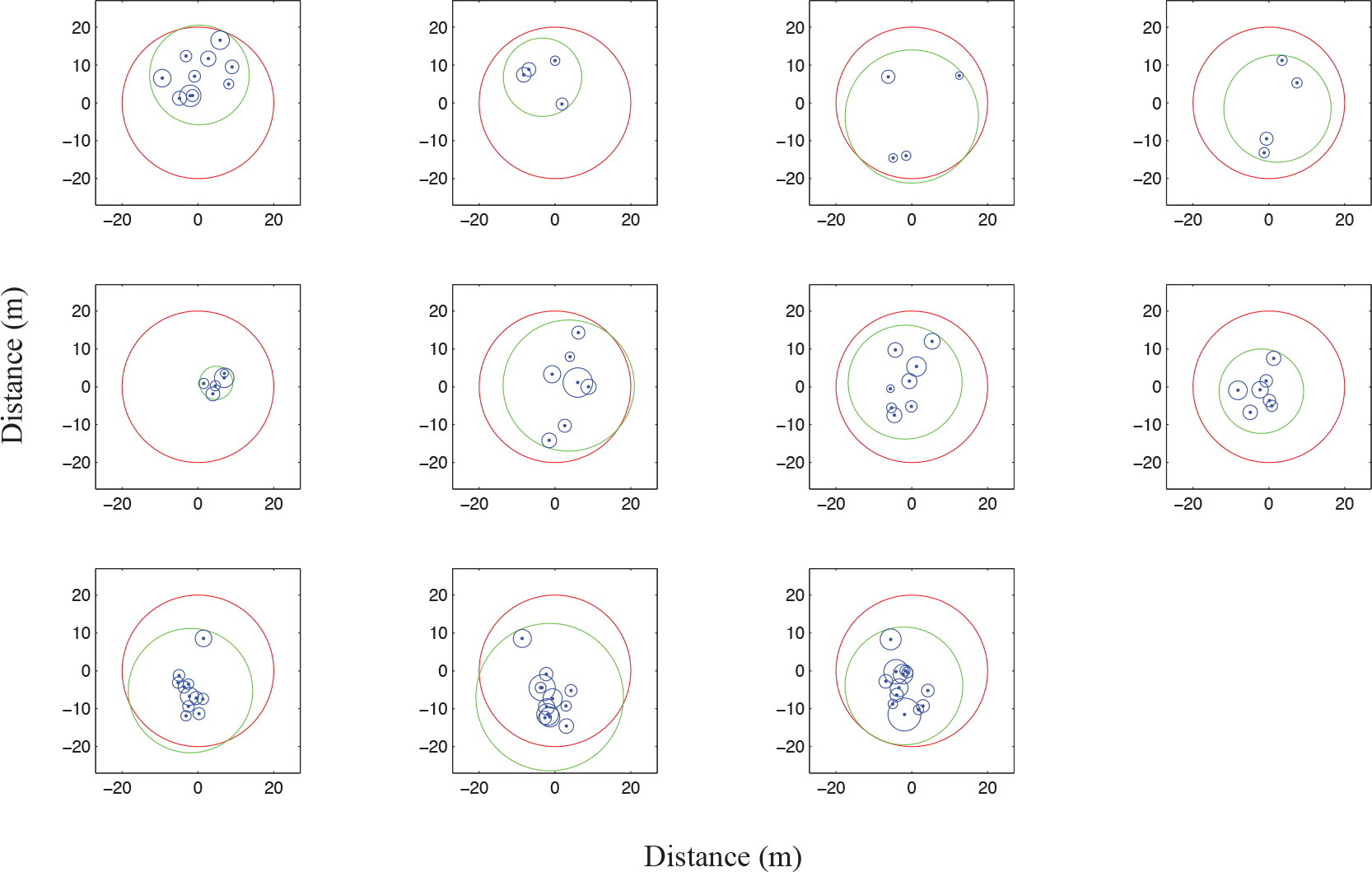
Chorus maps and sampling area. The red and the green circles represent ‘Large’ and ‘Small’ sampling area respectively for each of the 11 choruses. The blue dots and circles represent the position and broadcast area of callers.

**Figure A2.**
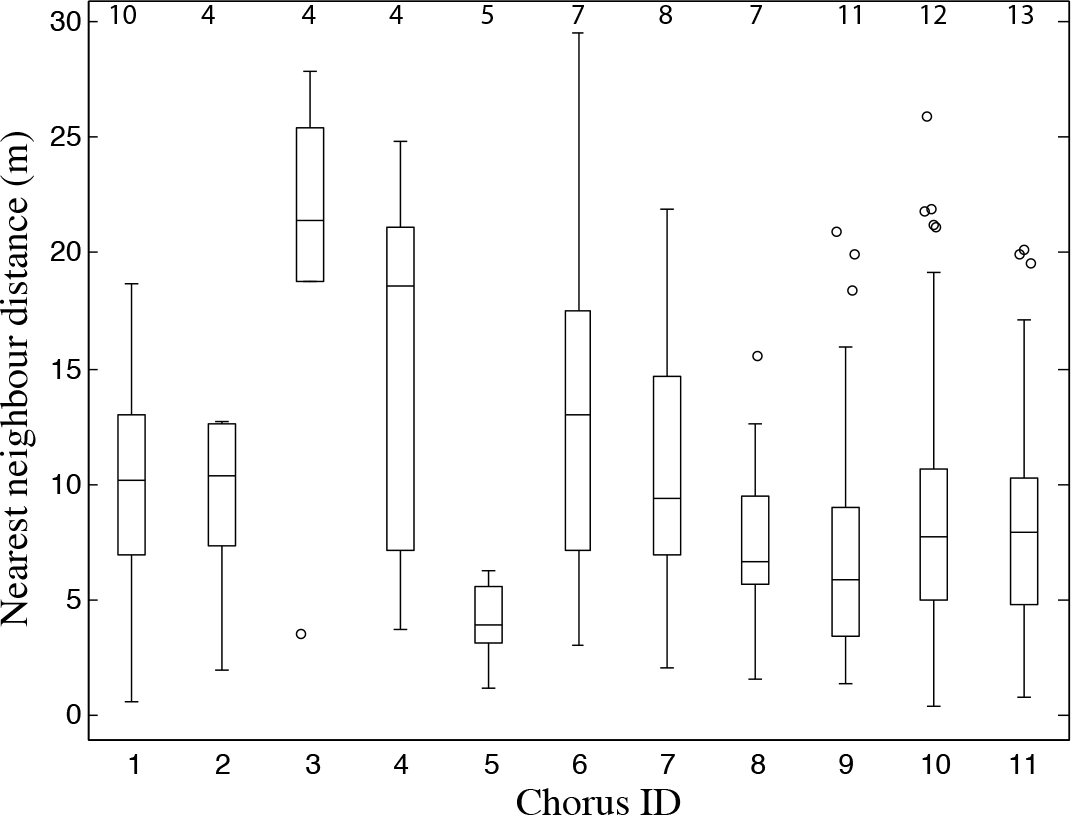
Distributions of nearest neighbor distances of the 11 choruses which were used in the simulations. The number at the top of each boxplot corresponds to the number of callers in the chorus.

**Figure A3:**
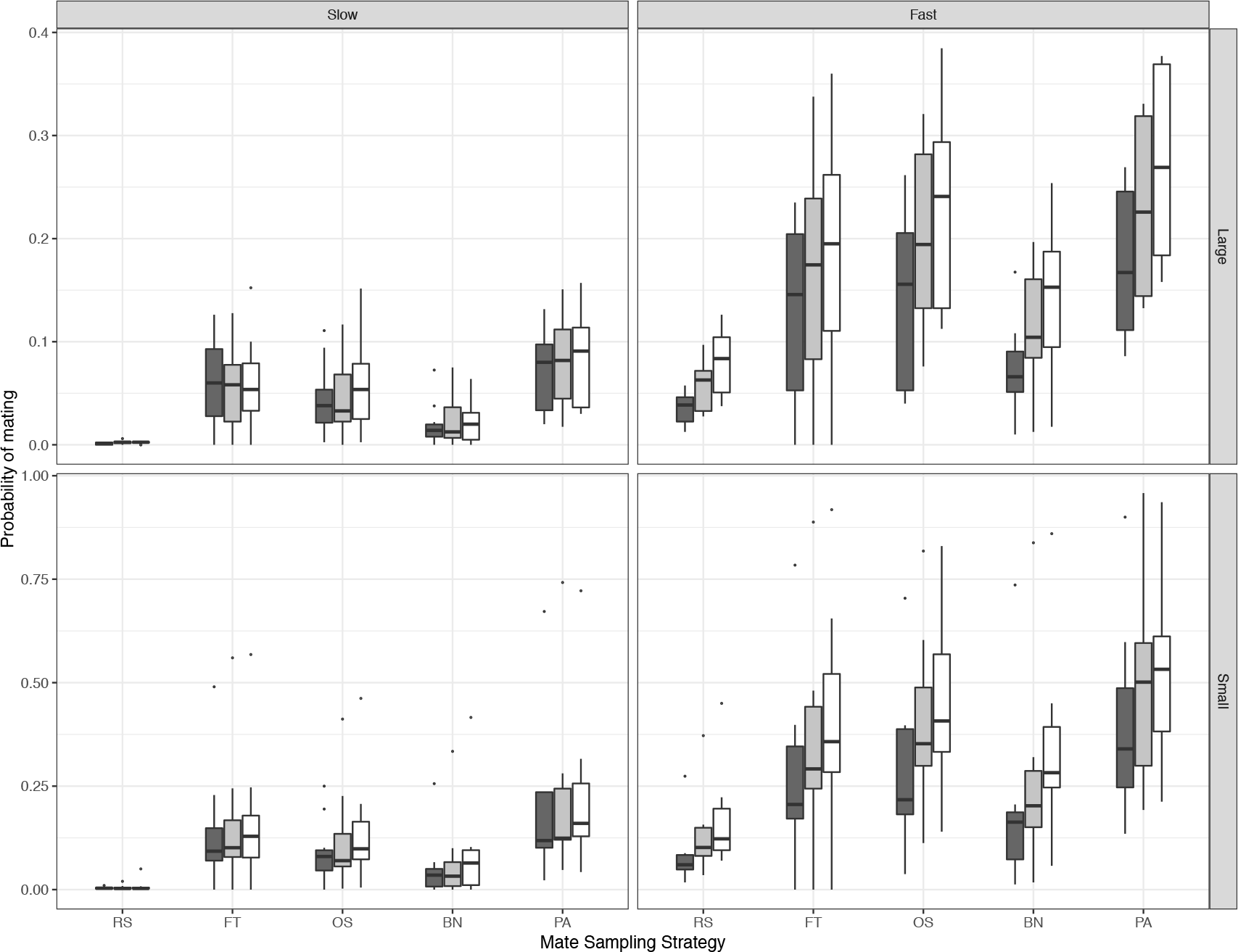
Probability of mating for different mate sampling strategies. The probability of mating for a female using each of the different mate sampling strategies across all the simulation sets. Grey = 3 hours, Light grey = 6 hours, White = 9 hours time window available for sampling. RS = Random sampling; FT = Fixed-threshold; OS = One-step decision, BN= Best-of-n, PA= Passive attraction. ‘Large’ = Large Area, ‘Small’= Small Area

**Figure A4:**
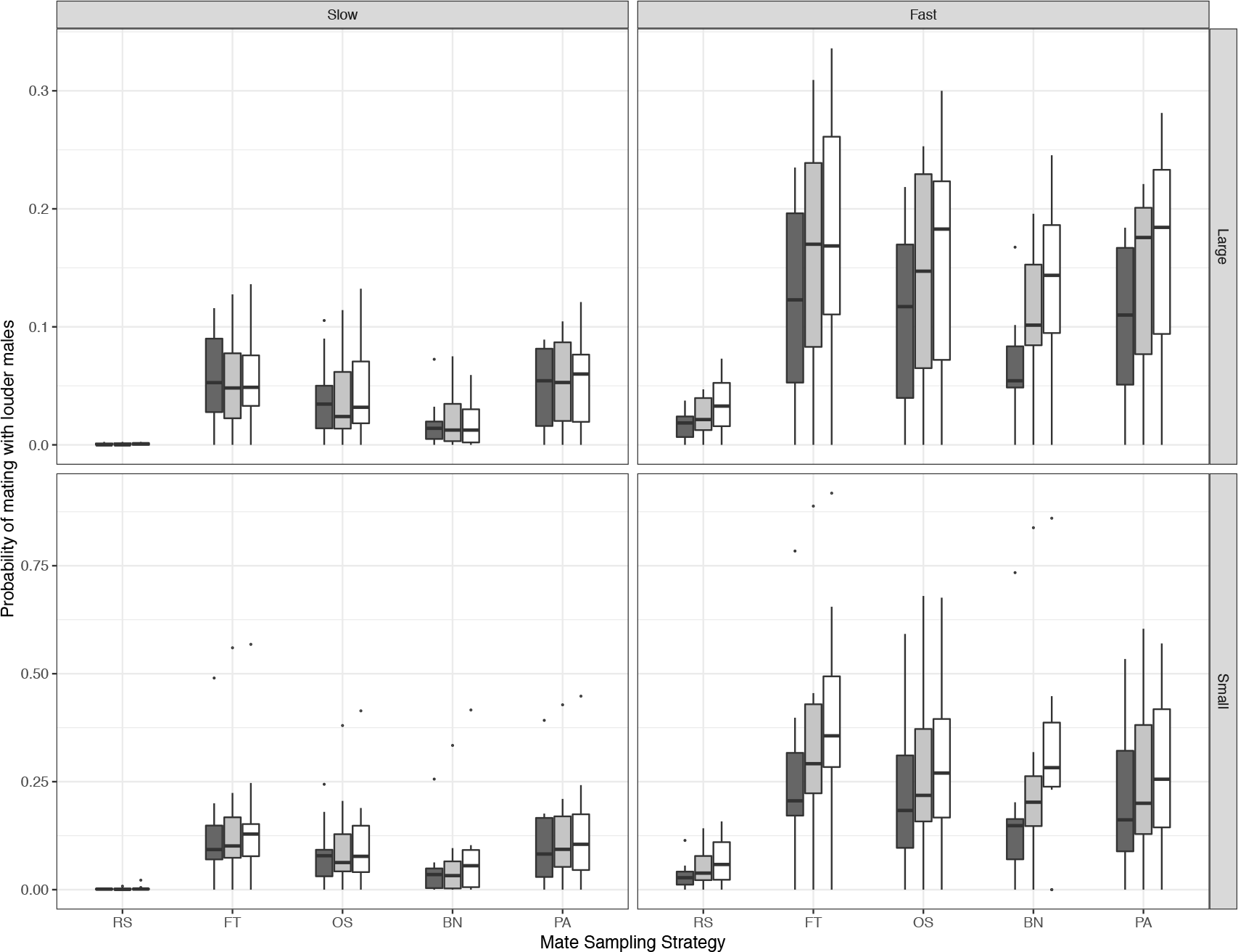
Probability of mating with a louder male for different mate sampling strategies. The probability of mating with a louder male for a female using each of the different mate sampling strategies across all the simulation sets. Grey = 3 hours, Light grey = 6 hours, White = 9 hours time window available for sampling. RS = Random sampling; FT = Fixed-threshold; OS = One-step decision, BN= Best-of-n, PA= Passive attraction. ‘Large’ = Large Area, ‘Small’=Small Area

**Figure A5:**
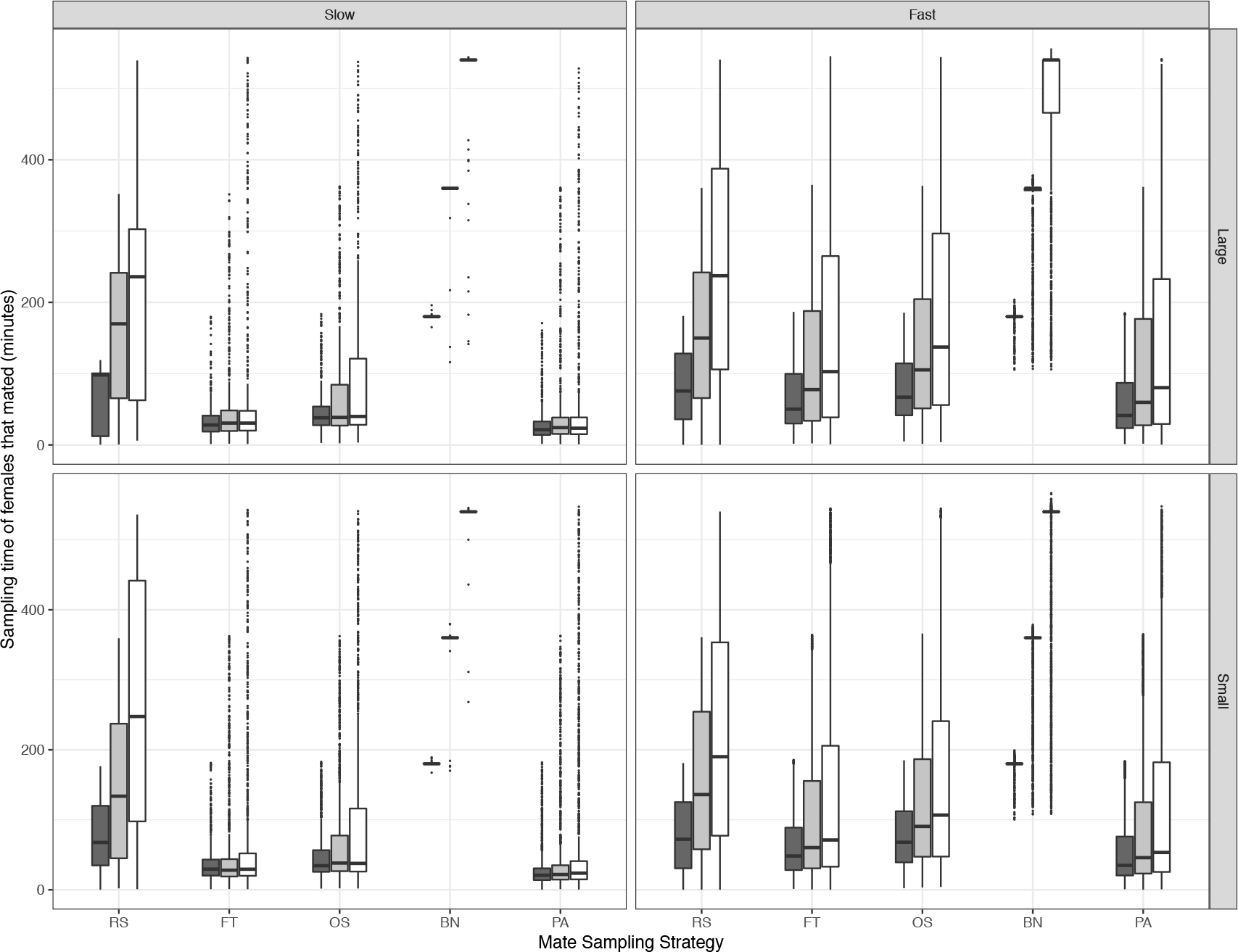
Costs of different sampling strategies. Distribution of sampling time of females successful in mating within the sampling window, using each of the five mate sampling strategies across all the simulation sets. Grey = 3 hours, Light grey = 6 hours, White = 9 hours time window available for sampling. ‘Large’ = Large Area, ‘Small’= Small Area

**Figure A6:**
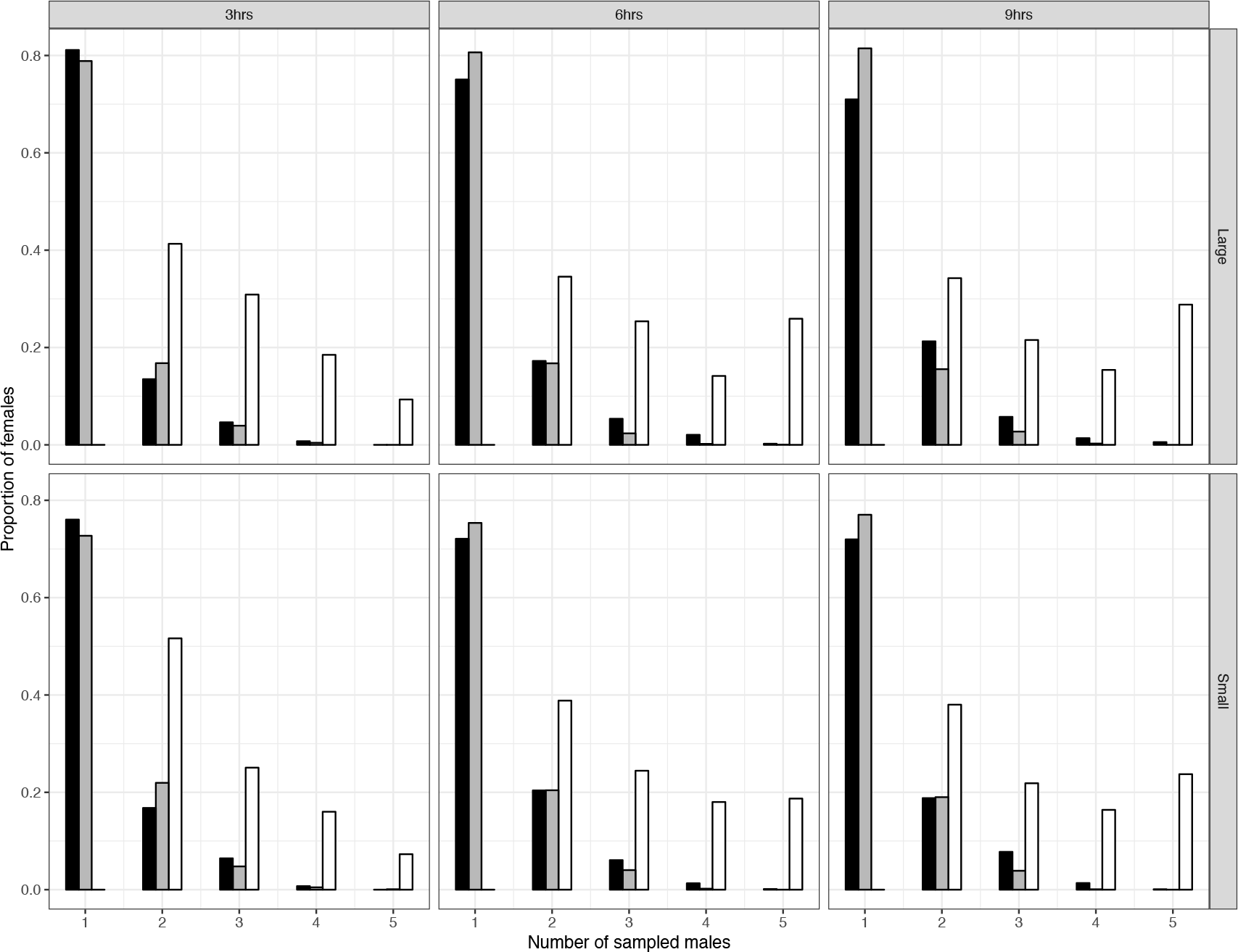
Number of sampled males. Relative frequency distribution of the number of males sampled by females that mated using each of the three active sampling strategies (FT = Fixed-threshold; OS = One-step decision, BN= Best-of-n), for ‘Fast’ females across the different simulation sets. Black = 3 hours, Grey = 6 hours, White = 9 hours of time window available for sampling. ‘Large’ = Large Area, ‘Small’= Small Area.

**Figure A7:**
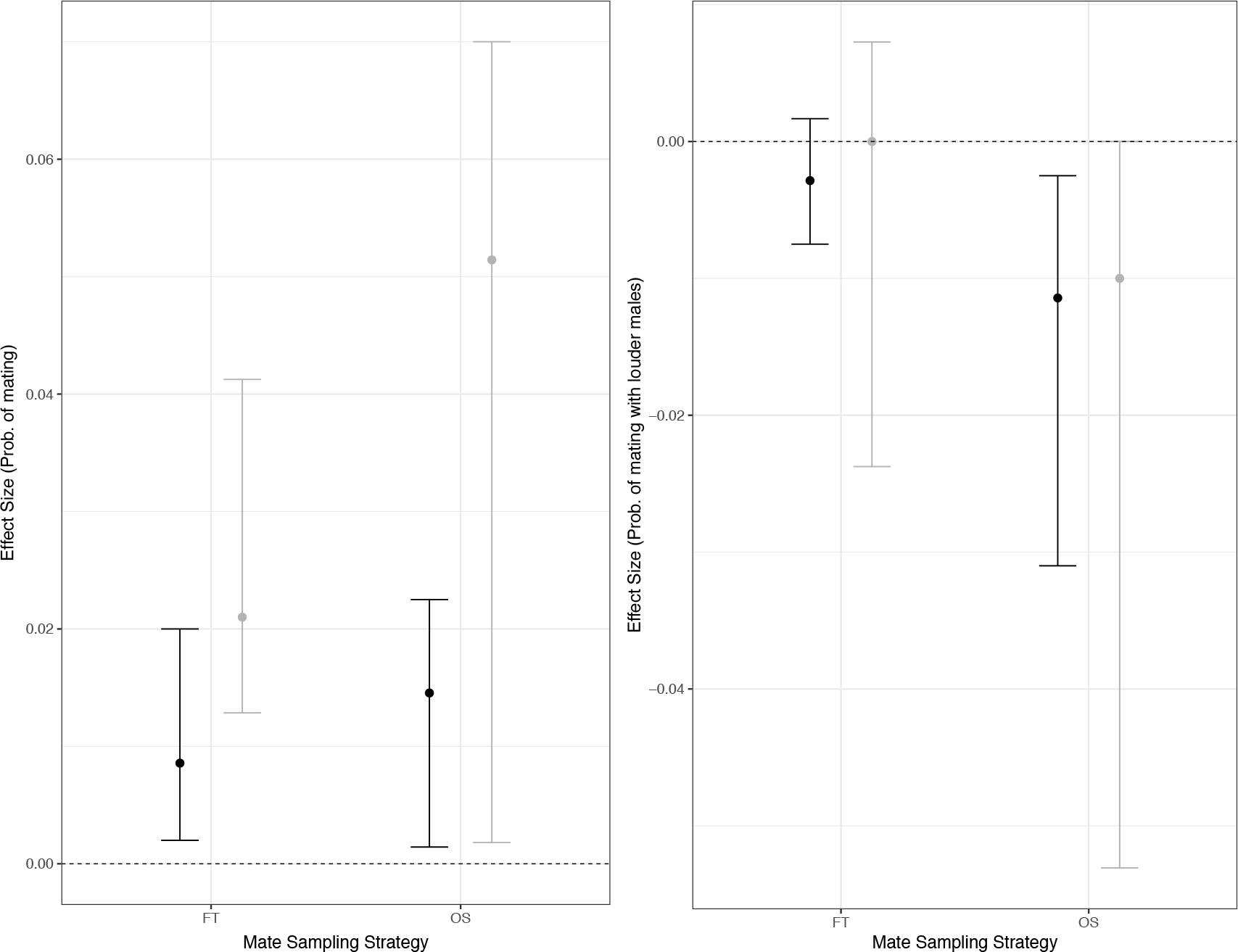
Effects of reducing the threshold criterion. The circles and the error-bars represent the medians and confidence intervals of the bootstrapped distributions of the median pairwise differences (choruswise) in the (a) probability of mating and (b) probability of mating with louder males between ‘passive attraction’ and the threshold strategies with reduced thresholds. The black and grey colours represent ‘Slow’ and ‘Fast’ females respectively. FT = Fixed-threshold; OS = One-step decision.

**Table A1.**
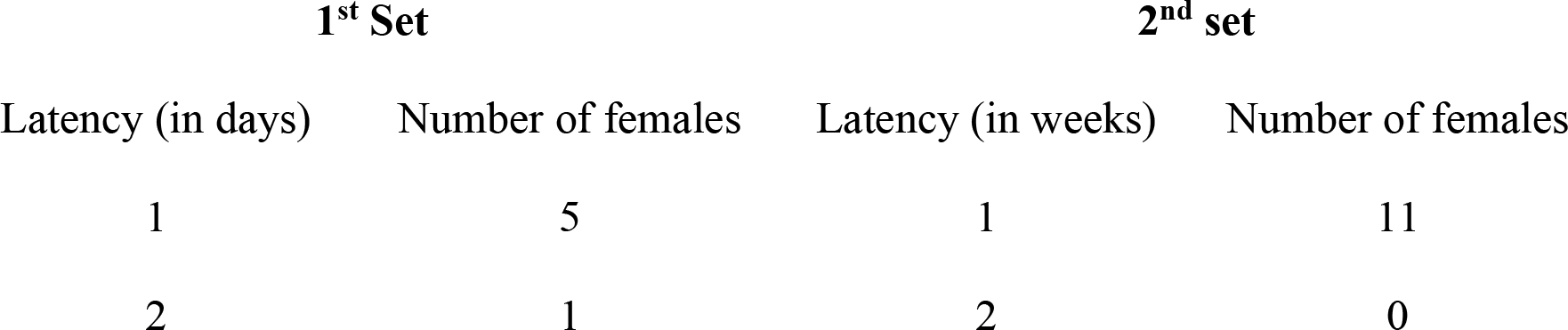
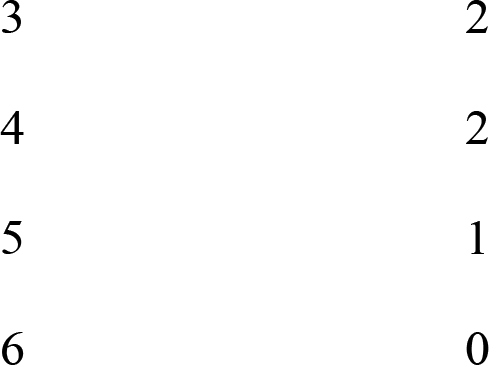
Latency of second matings across the two sets of remating experiments. In the first set 4 females were followed only for 5 days.

| 1<sup>st</sup> Set |  | 2<sup>nd</sup> set |  |
| --- | --- | --- | --- |
| Latency (in days) | Number of females | Latency (in weeks) | Number of females |
| 1 | 5 | 1 | 11 |
| 2 | 1 | 2 | 0 |

|  |  |
|---|---|
| 3 | 2 |
| 4 | 2 |
| 5 | 1 |
| 6 | 0 |

## References

Andersson, M. 1994. Sexual Selection. Princeton University Press, Princeton, New Jersey.

Bakker, T. C. M., and M. Milinski. 1991. Sequential female choice and the previous male effect in sticklebacks. Behavioral Ecology and Sociobiology 29:205–210.

Beckers, O. M., and W. E. Wagner Jr. 2011. Mate sampling strategy in a field cricket: evidence for a fixed threshold strategy with last chance option. Animal Behaviour 81:519–527.

Bensch, S., and D. Hasselquist. 1992. Evidence for active female choice in a polygynous warbler. Animal Behaviour 44, Part 2:301–311.

Berger-Tal, R., and Y. Lubin. 2011. High male mate search costs and a female-biased sex ratio shape the male mating strategy in a desert spider. Animal Behaviour 82:853–859.

Byers, J. A., P. A. Wiseman, L. Jones, and T. J. Roffe. 2005. A large cost of female mate sampling in pronghorn. The American Naturalist 166:661–668.

Castellano, S., and P. Cermelli. 2011. Sampling and assessment accuracy in mate choice: A random-walk model of information processing in mating decision. Journal of Theoretical Biology 274:161–169.

Castellano, S., A. Rosso, F. Laoretti, S. Doglio, and C. Giacoma. 2000. Call intensity and female preferences in the European green toad. Ethology 106:1129–1141.

Choudhury, S., and J. M. Black. 1993. Mate-selection behaviour and sampling strategies in geese. Animal Behaviour 46:747–757.

Dale, S., T. Amundsen, J. T. Lifjeld, and T. Slagsvold. 1990. Mate sampling behaviour of female pied flycatchers: evidence for active mate choice. Behavioral Ecology and Sociobiology 27:87–91.

Dale, S., H. Rinden, and T. Slagsvold. 1992. Competition for a mate restricts mate search of female pied flycatchers. Behavioral Ecology and Sociobiology 30:165–176.

Dombrovsky, Y., and N. Perrin. 1994. On Adaptive Search and Optimal Stopping in Sequential Mate Choice. The American Naturalist 144:355–361.

Forrest, T. G., and D. M. Green. 1991. Sexual selection and female choice in mole crickets (Scapteriscus: Gryllotalpidae): modelling the effects of intensity and male spacing. Bioacoustics 3:93–109.

Forrest, T. G., and R. Raspet. 1994. Models of female choice in acoustic communication. Behavioral Ecology 5:293–303.

Forsgren, E. 1997. Mate sampling in a population of sand gobies. Animal Behaviour 53:267–276.

Gerhardt, H. C., and F. Huber. 2002. Acoustic Communication in Insects and Anurans: Common Problems and Diverse Solutions. University of Chicago Press, Chicago and London.

Gibson, R. M., and T. A. Langen. 1996. How do animals choose their mates? Trends in Ecology & Evolution 11:468–470.

Hovi, M., and O. Rätti. 1994. Mate sampling and assessment procedures in female pied flycatchers (Ficedula hypoleuca). Ethology 96:127–137.

Janetos, A. C. 1980. Strategies of female mate choice: a theoretical analysis. Behavioral Ecology and Sociobiology 7:107–112.

Johnstone, R. A. 1997. The tactics of mutual mate choice and competitive search. Behavioral Ecology and Sociobiology 40:51–59.

Kasumovic, M. M., M. J. Bruce, M. E. Herberstein, and M. C. B. Andrade. 2007. Risky mate search and mate preference in the golden orb-web spider (Nephila plumipes). Behavioral Ecology 18:189–195.

Kokko, H., I. Booksmythe, and M. D. Jennions. 2014. Mate-sampling costs and sexy sons. Journal of Evolutionary Biology 28:259–266.

Kokko, H., M. D. Jennions, and R. Brooks. 2006. Unifying and testing models of sexual selection. Annual Review of Ecology, Evolution, and Systematics 37:43–66.

Kotiaho, J. S., and M. Puurtinen. 2007. Mate choice for indirect genetic benefits: scrutiny of the current paradigm. Functional Ecology 21:638–644.

Luttbeg, B. 1996. A comparative Bayes tactic for mate assessment and choice. Behavioral Ecology 7:451–460.

Meuche, I., O. Brusa, K. Linsenmair, A. Keller, and H. Pröhl. 2013. Only distance matters – non-choosy females in a poison frog population. Frontiers in Zoology 10:1:16.

Mhatre, N., and R. Balakrishnan. 2006. Male spacing behaviour and acoustic interactions in a field cricket: implications for female mate choice. Animal Behaviour 72:1045–1058.

Mhatre, N., and R. Balakrishnan. 2007. Phonotactic walking paths of field crickets in closed-loop conditions and their simulation using a stochastic model. Journal of Experimental Biology 210:3661–3676.

Mhatre, N., and R. Balakrishnan. 2008. Predicting acoustic orientation in complex real-world environments. Journal of Experimental Biology 211:2779–2785.

Milinski, M., and T. C. M. Bakker. 1992. Costs influence sequential mate choice in sticklebacks, Gasterosteus aculeatus. Proceedings of the Royal Society B 250:229–233.

Murphy, C. G., and H. C. Gerhardt. 2002. Mate sampling by female barking treefrogs (Hyla gratiosa). Behavioral Ecology 13:472–480.

Nakagawa, S., and I. C. Cuthill. 2007. Effect size, confidence interval and statistical significance: a practical guide for biologists. Biological Reviews 82:591–605.

Nandi, D. 2016. Acoustic signals, mate choice and mate sampling strategies in a field cricket (PhD). Indian Institute of Science, Bangalore, India.

Nandi, D., and R. Balakrishnan. 2013. Call intensity is a repeatable and dominant acoustic feature determining male call attractiveness in a field cricket. Animal Behaviour 86:1003–1012.

Nandi, D., and R. Balakrishnan. 2016. Spatio-Temporal Dynamics of Field Cricket Calling Behaviour: Implications for Female Mate Search and Mate Choice. PLOS ONE 11:e0165807.

Parker, G. 1982. Phenotype-limited evolutionarily stable strategies. Pages 173–202 in King’s college sociobiology group, ed. Current problems in sociobiology. Cambridge University Press, Cambridge, UK.

Parker, G. 1983. Mate quality and mating decisions. Pages 141–166 in P. Bateson, ed. Mate choice. Cambridge University Press, Cambridge, UK.

Petrie, M., H. Tim, and S. Carolyn. 1991. Peahens prefer peacocks with elaborate trains. Animal Behaviour 41:323–331.

R Development Core Team. 2014. R: A Language and Environment for Statistical Computing. R Foundation for Statistical Computing, Vienna, Austria.

Real, L. 1990. Search Theory and Mate Choice. I. Models of Single-Sex Discrimination. The American Naturalist 136:376–405.

Reid, M. L., and J. A. Stamps. 1997. Female mate choice tactics in a resource based mating system: field tests of alternative models. The American Naturalist 150:98–121.

Reinhold, K., and H. Schielzeth. 2014. Choosiness, a neglected aspect of preference functions: a review of methods, challenges and statistical approaches. Journal of Comparative Physiology A 201:171–182.

Rintamäki, P. T., R. V. Alatalo, J. Höglund, and A. Lundberg. 1995. Mate sampling behaviour of black grouse females (Tetrao tetrix). Behavioral Ecology and Sociobiology 37:209–215.

Ritz, M. S., and G. Köhler. 2007. Male behaviour over the season in a wild population of the field cricket Gryllus campestris L. Ecological Entomology 32:384–392.

Rodríguez-Muñoz, R., A. Bretman, J. Slate, C. A. Walling, and T. Tregenza. 2010. Natural and Sexual Selection in a Wild Insect Population. Science 328:1269–1272.

Rodríguez-Muñoz, R., A. Bretman, and T. Tregenza. 2011. Guarding Males Protect Females from Predation in a Wild Insect. Current Biology 21:1716–1719.

Roff, D. A., and D. J. Fairbairn. 2015. Bias in the heritability of preference and its potential impact on the evolution of mate choice. Heredity 114:404–412.

Stout, J. F., and R. McGhee. 1988. Attractiveness of the male Acheta domestica calling song to females. Journal of Comparative Physiology A 164:277–287.

Trail, P. W., and E. S. Adams. 1989. Active mate choice at cock-of-the-rock leks: tactics of sampling and comparison. Behavioral Ecology and Sociobiology 25:283–292.

Uy, J. A. C., L. Patricelli Gail, and G. Borgia. 2001. Complex Mate Searching in the Satin Bowerbird Ptilonorhynchus violaceus. The American Naturalist 158:530–542.

Wagner Jr., W. E., M. R. Smeds, and D. D. Wiegmann. 2001. Experience affects female responses to male song in the variable field cricket Gryllus lineaticeps (Orthoptera, Gryllidae). Ethology 107:769–776.

White, J. W., A. Rassweiler, J. F. Samhouri, A. C. Stier, and C. White. 2014. Ecologists should not use statistical significance tests to interpret simulation model results. Oikos 123:385–388.

Wickman, P.-O., and P. Jansson. 1997. An estimate of female mate searching costs in the lekking butterfly Coenonympha pamphilus. Behavioral Ecology and Sociobiology 40:321–328.

Wiegmann, D. D. 2000. Search behaviour and mate choice by female field crickets, Gryllus integer. Animal behaviour 58:1293–1298.

Wiegmann, D. D., and L. M. Angeloni. 2007. Mate choice and uncertainty in the decision process. Journal of Theoretical Biology 249:654–666.

Wiegmann, D. D., L. A. Real, T. A. Capone, and S. Ellner. 1996. Some distinguishing features of models of search behavior and mate choice. American Naturalist 147:188–204.

Wiegmann, D. D., S. M. Seubert, and G. A. Wade. 2010. Mate choice and optimal search behavior: Fitness returns under the fixed sample and sequential search strategies. Journal of Theoretical Biology 262:596–600.

Wittenberger, J. 1983. Tactics of mate choice. Pages 435–447 in P. Bateson, ed. Mate choice. Cambridge University Press, Cambridge, UK.

